# Engineering morphology of tetrasulfide-incorporated mesoporous silica nanoparticles for enhanced mRNA delivery

**DOI:** 10.64898/2026.11.03.763884

**Authors:** Zhe Cao, Silja Saarela, Jiajia Wang, Minna Sivonen, Emilia Happonen, Jussi Hepojoki, Hiromi Muramatsu, Pardi Norbert, Vesa-Pekka Lehto, Wujun Xu

**Affiliations:** Department of Technical Physics, University of Eastern Finland, Kuopio 70211, Finland; Department of Virology, Faculty of Medicine, University of Helsinki, Finland; Department of Microbiology, Perelman School of Medicine, University of Pennsylvania, Philadelphia, PA, USA

**Keywords:** Virus-like mesoporous silica, Tetrasulfide, Lipid nanoparticles, mRNA delivery, Nanospikes

## Abstract

Lipid nanoparticles (LNPs) are the most studied nanocarrier for mRNA delivery, yet their broader application remains limited due to suboptimal cellular uptake, low efficacy in mRNA delivery, and poor stability. In this study, we report a novel tetrasulfide-incorporated virus-like porous silica (VLPSi) nanoparticle as a safe and efficient platform to enhance LNP-mediated mRNA delivery. By systematically tuning the synthesis parameters, a series of nanoparticles with distinct morphologies and architectures were obtained, enabling the study of how tetrasulfide and particle morphology affect mRNA delivery. The performance of VLPSi was optimized by evaluating cellular uptake, cytotoxicity, and transfection performance with different lipid and mRNA payloads across multiple cell types. The results showed that increasing the length and number of spikes on the VLPSi surface significantly improved cellular uptake efficiency and subsequent mRNA expression. Mechanistically, the enhanced delivery arose from the combined effect from the nanospikes and intracellular tetrasulfide degradation. Furthermore, the nanoparticles significantly enhanced the stability of mRNA. Overall, these findings establish a structure-function relationship between virus-like topography, biodegradable silica chemistry, and mRNA delivery. This work provides a structurally tunable, biodegradable, and highly effective bioinspired nanoplatform to enhance mRNA delivery.

## 1. Introduction

Messenger RNA (mRNA)-based therapeutics have gained increasing attention in vaccines developments, protein replacement therapy, and cancer immunotherapy due to its advantages such as transient expression and no known risk of genome integration^1,2^. However, naked mRNA is easily degraded by enzymes and has often relatively low cellular uptake efficiency^3^. It also faces challenges in crossing multiple cellular barriers, including cell membrane and endosomal membrane^4^. Therefore, the development of safe and efficient mRNA delivery systems is crucial.

Currently, the development of mRNA delivery systems mainly focuses on lipid nanoparticles (LNPs). LNPs usually contain 4 different lipids, including the ionizable lipid, helper phospholipid, cholesterol and PEG-lipid. The ionizable lipid becomes protonated in acidic endosomal environments, thereby disrupting endosomal structures allowing cytoplasmic delivery of the mRNA cargo^5^. LNPs can efficiently encapsulate mRNA, preventing it from enzyme-degradation during extracellular transport^6^. However, the mRNA delivery efficacy of conventional LNPs is low due to the low efficiency of cellular uptake and endosomal escape. Furthermore, mRNA-LNPs typically have poor thermal stability^7^. These limitations have prompted the exploration of new carrier platforms with enhanced cellular uptake, mRNA delivery, and stability.

Mesoporous silica nanoparticles have attracted widespread attention in the biomedical field due to their high surface area, tunable pore size, low toxicity, and controllable chemical properties^8^. In particular, virus-like mesoporous silica (VLPSi) nanoparticles with a spiked surface morphology exhibit higher cellular uptake as compared to spherical particles^9–11^. Recently, we demonstrated VLPSi showed a great promise to enhance the delivery of mRNA^12,13^. However, the VLPSi is composed of Si-O-Si structures, which are relatively stable inside cells and may hinder the effective release of their loads^14^. Current research indicates that incorporating tetrasulfide bonds can enable silica nanoparticles to specifically respond to the high GSH environment within cells, leading to rapid degradation^15,16^. However, the influence of tetrasulfide bond incorporation on the formation mechanism and morphological change of VLPSi has not been systematically studied. Furthermore, current research mostly focuses on integrating tetra sulfide bonds into porous silica for drug delivery while the research on mRNA delivery is lacking^17^.

The present study aims to investigate how the incorporation of tetrasulfide bonds into VLPSi affects particle size and morphology. Based on this, we further investigated the influence of different experimental parameters on the VLPSi structure. Finally, we studied the influences of these alterations to mRNA delivery efficiency. The results showed that the incorporation of tetrasulfide bonds significantly affected the structure of the nanoparticles, especially the length and number of spikes. VLPSi-S5 with the highest spike-density and longest spikes can achieve the best mRNA transfection efficiency as judged by protein expression in vitro. This indicates that the virus-like structure can significantly improve cellular uptake, and that the transfection efficacy is further enhanced by increasing spike length and density. Furthermore, the incorporation of tetrasulfide bonds ensures the intracellular degradation of nanoparticles, promoting mRNA release from nanoparticles and thus improving transfection. The redox reaction of the tetrasulfide bonds can further modulate intracellular GSH levels and promote mRNA translation (Scheme 1). In summary, the present study provides insights into constructing novel mRNA transfection platforms and lay the foundation for future development of efficient mRNA transfection methods.

**Scheme 1.**
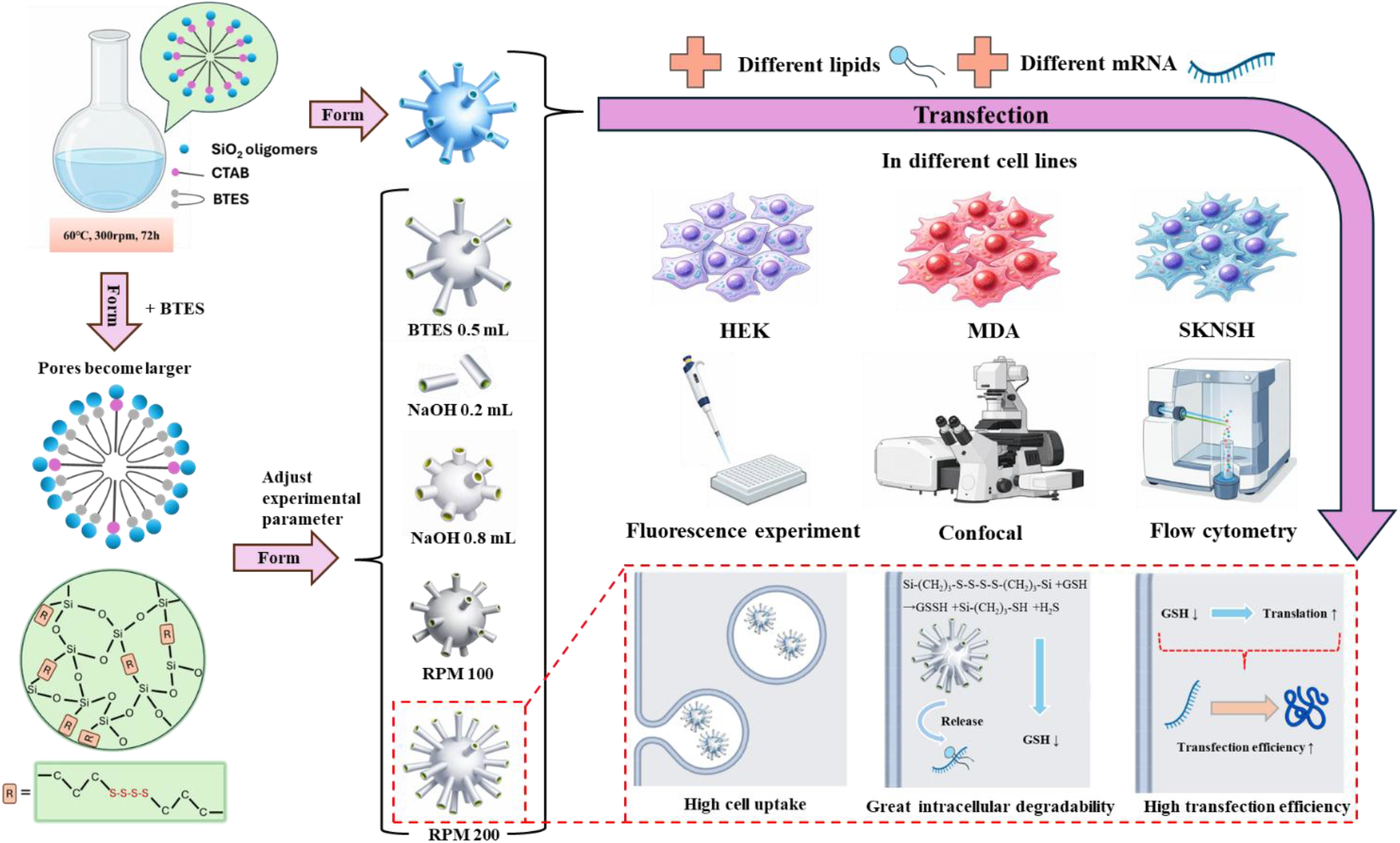
Synthesis of different nanoparticles and their application in mRNA transfection

## 2. Materials and methods

### 2.1. Synthesis of VLPSi and VLPSi-S

a) VLPSi: This method was adapted from Wang et al. and modified accordingly^18^. Hexadecyl trimethyl ammonium Bromide (CTAB, Sigma) was dissolved in 40 mL deionized water and heated in an oil bath at 60 °C for 15 min. 0.4 mL of 0.1 M NaOH was dissolved in the above aqueous solution. Subsequently, 4 mL of tetraethyl orthosilicate (TEOS, Merck) and 16 mL of cyclohexane (Merck) were mixed thoroughly and added dropwise to the above solution. The mixture was stirred continuously at 60 °C and 300 rpm for 72 h. The product was collected by centrifugation and washed several times with water and ethanol.

b) VLPSi-S: CTAB was dissolved in 40 mL deionized water and heated in an oil bath at 60 °C for 15 minutes. 0.4 mL of 0.1 M NaOH was dissolved in the above aqueous solution. Subsequently, 4 mL of TEOS, 0.5 mL of organosilane precursor Bis[3-(triethoxysilyl) propyl] tetrasulfide (BTES, Sigma), and 16 mL of cyclohexane were mixed thoroughly and added dropwise to the above solution. The mixture was stirred continuously at 60 °C and 300 rpm for 72 h. The product was collected by centrifugation and washed several times with water and ethanol.

The methods for synthesizing other nanoparticles are based on VLPSi-S1 with modified experimental parameters, as shown in the table below:

**Table S1.** Experimental conditions for different nanoparticles.

| Nanoparticles | TEOS (mL) | BTES (mL) | NaOH (mL) | Stirring rate (rpm) |
|---|---|---|---|---|
| VLPSi-S1 | 4 | 0.5 | 0.4 | 300 |
| VLPSi-S2 | 4 | 0.5 | 0.2 | 300 |
| VLPSi-S3 | 4 | 0.5 | 0.8 | 300 |
| VLPSi-S4 | 4 | 0.5 | 0.4 | 100 |
| VLPSi-S5 | 4 | 0.5 | 0.4 | 200 |
| VLPSi-S6 | 4 | 1 | 0.4 | 300 |
| VLPSi-S7 | 4 | 2 | 0.4 | 300 |

All nanoparticles were refluxed at 78°C for 72 h with an ethanol solution (10% V/V) containing hydrochloric acid (HCl) to completely remove the CTAB template.

### 2.2. Characterization of nanoparticles

The nanoparticles morphology was characterized by using transmission electron microscopy (TEM) (Jeol JEM-2100F, Japan). ImageJ was used to analyze TEM images of different nanoparticles to measure the diameter, core size, spike length and number of the nanoparticles (particle number n = 100 per sample). The structure and chemical composition of different nanoparticles were determined using Fourier Transform Infrared Spectroscopy (FTIR) (Thermo Nicolet iS50, USA). The elemental composition of different nanoparticles was determined using X-ray photoelectron spectroscopy (XPS) (Thermo Nexsa G2, USA). The zeta potential of different nanoparticles was measured by Zetasizer Nano ZS (Malvern Panalytical Ltd, UK).

### 2.3. Degradability test of VLPSi and VLPSi-S

L-Glutathione reduced (GSH, Merck) was dissolved in deionized water to prepare GSH solutions with concentrations of 5 μM, 5 mM, and 10 mM to simulate different physiological environments. 100 μg/mL of samples were placed in GSH solutions of different concentrations and sonicated thoroughly, then incubated with shaking in an incubator at 37°C. Samples were taken at different time points (0, 8, 16, 24h) for TEM to observe the degradation of nanoparticles.

### 2.4. Cell culture

Human embryonic kidney cells (HEK-293T) were cultured in Dulbecco’s Modified Eagle Medium (DMEM) at 37 °C and 5% CO_2_, supplemented with 10% fetal bovine serum (FBS, Sigma) and 1% penicillin-streptomycin (Gibco). When the HEK-293T cells in flask reach about 80-90%, wash three times with phosphate-buffered saline (PBS) and detached with 0.25% Trypsin-EDTA (Gibco) for passage. Human triple negative breast cancer cells, MDA-MB-231 (MDA) were cultured in Roswell Park Memorial Institute (RPMI-1640) medium at 37°C and 5% CO_2_, supplemented with 10% fetal bovine serum (FBS, Sigma) and 1% penicillin-streptomycin (Gibco). The human neuroblastoma cell (SK-N-SH) were cultured and passaged using the same methods and medium as HEK-293T cells.

### 2.5. Cell cytotoxicity

50 μL of Poly-L-Lysine (PLL) was coated on each well of a 96-well plate beforehand to facilitate HEK-293T cell attachment. Cells were then cultured for 24 h, with cell number is 10⁴ cells per well. Different nanoparticles were then added to the wells for further cultivation, with final concentrations of 5, 10, 20, 40, 80, and 160 μg/mL per well. The treatment without nanoparticles served as a control. After incubation at 37℃ for 24 h, the culture medium containing nanoparticles was removed, and the plates were washed three times with PBS. Then, 50 μL medium and 50 μL CellTiter-Glo luminescent detection reagent (Promega) was added in each well. The 96-well plate was placed in dark environment at room temperature for 15 min. Fluoroskan Ascent FL microplate reader (Thermo Labsystems) was used to read the fluorescence results. For relative quantitative analysis, the mean value of the control check (CK) was set as 100%, and data from other experimental groups were normalized to the CK values. The results are expressed as relative values.

### 2.6. Cellular uptake

HEK-293T cells were cultured in 8-well plates with 2 × 10⁴ cells per well and incubated at 37°C for 24 h. Different nanoparticles (VLPSi, VLPSi-S1, VLPSi-S2, VLPSi-S3, VLPSi-S4, VLPSi-S5) were selected for cell uptake experiments. First, amino groups were modified on the surface of different nanoparticles to facilitate subsequent binding with fluorescein 5(6)-isothiocyanate (FITC, Alfa Chemical)^19^. 10 mg of the modified nanoparticles were mixed with 1 mg FITC in 2 mL ethanol and stirred in dark at room temperature for 12 h. Subsequently, the FITC-labeled nanoparticles were collected by centrifugation and washed three times with ethanol. The labeled nanoparticles were added to 8-well plates at 40 μg/mL concentration per well and incubated at 37°C for 4 h. Excess particles were then washed away with PBS. The cells were stained with 5 μg/mL CellMask™ Plasma Membrane stains (Invitrogen, USA) and incubated in an incubator for 10 min. They were then washed three times with PBS and fixed with 4% formaldehyde (Sigma-Aldrich) at room temperature for 10 min. Staining was performed for 10 min using 10 μg/mL DAP1 (4′, 6-diamide-2-benzodiazepine, Sigma-Aldrich), followed by washing three times with PBS. Finally, 200 μL PBS was added to each well. Images were captured using a fluorescence imaging confocal laser scanning microscope (Zeiss LSM 800) and processed using ZEISS ZEN software.

### 2.7. mRNA production

Firefly luciferase (Luc)-encoding mRNA (Luc mRNA) was synthesized as described earlier^20^. Briefly, the codon-optimized DNA sequence for Luc was synthesized (Genscript) and cloned into a T7 promoter-driven mRNA production plasmid. The plasmid was linearized and subjected to *in vitro* transcription using the MEGAscript T7 kit (Ambion) to generate mRNAs with 101 nucleotide-long poly(A) tails. Co-transcriptional capping was performed using the CleanCap cap1 analog (TriLink), and m¹Ψ-5’-triphosphate (TriLink) was incorporated in place of uridine-triphosphate (UTP). Following transcription, mRNAs were purified using a cellulose-based method and analyzed by agarose gel electrophoresis before storage at −20 °C^20^.

mCherry mRNA was generated via transcription using plasmid DNA (pDNA) containing the mCherry sequence, provided by the laboratory of Jussi Hepojoki. Using this pDNA as a template, polymerase chain reaction (PCR) was performed to amplify the mCherry gene containing a T7 promoter. Specific procedures followed previous studies^21^, and the PCR reaction utilized Q5 High-Fidelity 2x Master Mix (New England BioLabs, USA). The amplification products were purified using the NucleoSpin Gel and PCR Clean-up kit (Macherey-Nagel, Germany). Subsequently, using purified DNA as a template, in vitro transcription and purification of mCherry mRNA were performed using the HiScribe T7 ARCA mRNA kit (New England BioLabs, USA). Next, the mRNA undergoes poly-A tailing. Finally, the concentration of mCherry mRNA dissolved in RNase-free water was determined using NanoDrop One UV-Vis spectrophotometer (Thermo Fisher Scientific, USA), and the final product was stored at −80 °C.

### 2.8. Flow cytometry analysis of mRNA delivery

#### 2.8.1. mCherry mRNA delivery effect of different nanoparticles with lipid

HEK-293T Cells were cultured in PLL-coated 24-well plates for 24 hours, with a cell number of 6 × 10⁴ cells per well. Based on the report of Meng et al., better mRNA delivery can be achieved by first binding mRNA to cationic polymers and then binding them to the surface of nanoparticles^22^. Following this assembly sequence, we first combined mCherry-encoding mRNA with a complex of an ionizable cationic lipid ALC-0315 mixed with a helper lipid DOPC (mass ratio ALC-0315/DOPC = 5:3) in an acidic environment. Then, we combined them with different nanoparticles (VLPSi, VLPSi-S1, VLPSi-S2, VLPSi-S3, VLPSi-S4, VLPSi-S5) to form a final complex and dissolve them in DMEM. Then the final complex was added in 24-well plate. The only culture medium was used as negative control, and Lipofectamine 2000 (Lipo, Thermo Fisher) was used as positive control. Each well contained 0.15 μg mCherry mRNA and 380 μL medium. The mass ratio of mCherry mRNA: Lipo = 1:4 and mCherry mRNA: ALC/DOPC: nanoparticles = 1:8:25. After incubation at 37°C for 4 hours, the old medium was replaced with DMEM (10% FBS). Incubation continued at 37°C for another 24 hours. The medium was then removed, and cells were incubated with Typsin and collected. After washing, FLOW buffer was added to disperse the cells evenly, and the cells were placed on ice for testing. The mCherry mRNA transfection efficiency of each group was determined using a CytoFLEX flow cytometer (Beckman Coulter, USA), and the data were analyzed using CytExpert v2 software (Beckman Coulter, USA).

#### 2.8.2. mCherry mRNA delivery effect of nanoparticles with different lipids

The same assembly sequence was used as in section 2.8.1, the final complexes of mCherry mRNA, different lipid (the mass ratio of ALC-0315/DOPC=5: 3, DOTAP/DOPC=2:1 and DOTMA/DOPC=2:1), and VLPSi-S5 were prepared and dissolved in DMEM. Next, prepare complexes of mCherry mRNA and different lipids without nanoparticles for comparison. Then the final complex was added in 24-well plate. The only culture medium was used as negative control, and Lipo was used as positive control. Each well contained 0.15 μg mCherry mRNA and 380 μL medium. The mass ratio of mCherry mRNA: Lipo = 1:4. mCherry mRNA: different lipids: VLPSi-S5 = 1:8:25. mCherry mRNA: different lipids=1:8. The remaining experimental parameters and procedures are the same as in section 2.8.1.

#### 2.8.3. mCherry mRNA delivery effect of different nanoparticles with lipid in different cell lines

HEK-293T, MDA and SKNSH cells were selected for the experiment. Using the same assembly sequence as in section 2.8.1, the final complexes of mCherry mRNA, ALC-0315/DOPC, and different nanoparticles (VLPSi, VLPSi-S4, VLPSi-S5) were prepared and dissolved in different mediums. Then the final complex was added in 24-well plate. The remaining experimental parameters and procedures are the same as in section 2.8.1.

### 2.9. Luciferase expression analysis after mRNA delivery

#### 2.9.1. Luc mRNA delivery with different nanoparticles loaded with lipid

HEK-293T cells were cultured in PLL-coated 96-well plates for 24 hours with 2 × 10⁴ cells per well. Using the same assembly sequence as in section 2.8.1, the final complexes of Luc mRNA, ALC-0315/DOPC, and different nanoparticles (VLPSi, VLPSi-S1, VLPSi-S2, VLPSi-S3, VLPSi-S4, VLPSi-S5) were prepared and dissolved in DMEM. Then the final complex was added in 96-well plate. The only culture medium was used as negative control, and Lipo was used as positive control. Each well contains 0.05 μg Luc mRNA and 110 μL medium. The mass ratio of Luc mRNA: Lipo = 1:4 and Luc mRNA: ALC-0315/DOPC: nanoparticles = 1:8:25. After incubation at 37°C for 4 hours, the old medium was replaced with DMEM (10% FBS). After further incubation at 37°C for 24 hours, 100 μL of ONE-Glo Luciferase assay reagent (Promega) was added to each well. After standing in the dark for 5 minutes, the fluorescence results were read using a Fluoroskan Ascent FL microplate reader. For relative quantitative analysis, the average value of the Lipo results was set to 1, and the data of other experimental groups were normalized according to CK and expressed as relative values.

#### 2.9.2. Luc mRNA delivery with nanoparticle loaded with different lipids

HEK-293T cells were selected for the experiment. Using the same assembly sequence as in section 2.8.1, the final complexes of Luc mRNA, different lipid (the mass ratio of ALC-0315/DOPC=5: 3, DOTAP/DOPC=2:1 and DOTMA/DOPC=2:1), and VLPSi-S5 were prepared and dissolved in DMEM. Next, prepare complexes of Luc mRNA and different lipids without nanoparticles for comparison. Then the final complex was added in 96-well plate. The only culture medium was used as negative control, and Lipo was used as positive control. Each well contained 0.05 μg Luc mRNA and 110 μL medium. The mass ratio of Luc mRNA: Lipo = 1:4. Luc mRNA: different lipids: VLPSi-S5 = 1:8:25. Luc mRNA: different lipids=1:8. The remaining experimental parameters and procedures are the same as in section 2.8.1.

#### 2.9.3. Luc mRNA delivery with different nanoparticles loaded with lipid in different cell lines

HEK-293T, MDA and SKNSH cells were selected for the experiment. Using the same assembly sequence as in section 2.8.1, the final complexes of Luc mRNA, ALC-0315/DOPC, and different nanoparticles (VLPSi, VLPSi-S4, VLPSi-S5) were prepared and dissolved in different mediums. Then the final complex was added in 96-well plate. The remaining experimental parameters and procedures are the same as in section 2.8.1.

### 2.10. Confocal microscopy analysis of mCherry mRNA transfection

HEK-293T cells were cultured in PLL-coated 96-well plates for 24 hours with 1.5 × 10⁴ cells per well. Using the same assembly sequence as in section 2.8.1, the final complexes of mCherry mRNA, ALC/DOPC, and different nanoparticles (VLPSi, VLPSi-S1, VLPSi-S2, VLPSi-S3, VLPSi-S4, VLPSi-S5) were prepared and dissolved in DMEM. Then the final complex was added in 96-well plate. The only culture medium was used as negative control, and Lipo was used as positive control. Each well contained 0.05 μg mCherry mRNA and 110 μL medium. mCherry mRNA: ALC/DOPC: nanoparticles = 1:8:25. After incubation at 37°C for 4 hours, the old medium was replaced with DMEM (10% FBS). After incubation at 37°C for another 24 hours, the old medium was removed and the sample was washed three times with PBS. Staining was performed using nuclear stain (DAPI) at a working concentration of 10 μg/mL. After staining for 10 min at room temperature, the samples were washed three times with PBS, then100 μL PBS was added to each well. Images were captured using a fluorescence imaging confocal laser scanning microscope (Zeiss LSM 800) and the image data were processed using ZEISS ZEN software.

### 2.11. Stability study

HEK-293T cells were selected for the experiment. Using the same assembly sequence as in section 2.8.1, the final complexes of Luc mRNA, ALC/DOPC, and different nanoparticles (VLPSi, VLPSi-S4, VLPSi-S5) were prepared and dissolved in different medias. They were then stored at 4°C for 7 days. 50 μL PLL was added to each well of 96-well plate. All PLL was removed before adding cells for culture. Cells were then cultured in 96-well plates for 24 hours, with a cell number of 2 × 10⁴ cells per well. The only culture medium was used as negative control, and Lipo was used as positive control. Subsequently, the complex prepared 7 days before, and the freshly complex using the same procedure as above was prepared and dissolved in mediums. Then the final complex was added in 96-well plate. All other experimental parameters and procedures were the same as in section 2.8.1.

### 2.12. Statistical Analysis

Statistical analysis was performed using Origin software. FlowJo 10.8.1 was used for the analysis and processing of flow cytometry data. ImageJ was used to statistically analyze the relevant parameters of nanoparticles and the fluorescence intensity of confocal images. Student’s T-test was used for comparative analysis to assess whether differences between groups were significant. Each experimental group contained at least three independent samples to ensure the stability and reproducibility of the results. Statistical significance is denoted as follows: \**p* < 0.05, \*\**p* < 0.01, \*\*\**p* < 0.001.

## 3. Result and discussion

### 3.1. Synthesis and Characterization of Nanoparticles

Spiky VLPSi with pure silicon dioxide (SiO_2_) was synthesized as previously^18^. To modify the nanoparticle with tetrasulfide bonds for enhanced biodegradability, Bis[3-(triethoxysilyl)propyl] tetrasulfide (BTES) with thioether bridging group was mixed with tetraethyl orthosilicate (TEOS) as silicon source for the synthesis of different VLPSi doped with tetrasulfide bonds (VLPSi-S). Subsequently, we investigated the effects of different reaction conditions on the morphology of nanoparticles, including BTES content, NaOH content, and experimental stirring rate (Table S1). The structures of the various nanoparticles are shown in Figure 1a, and the statistical data of nanoparticles are presented in Figure 1b, 1c and 1d.

**Figure 1.**
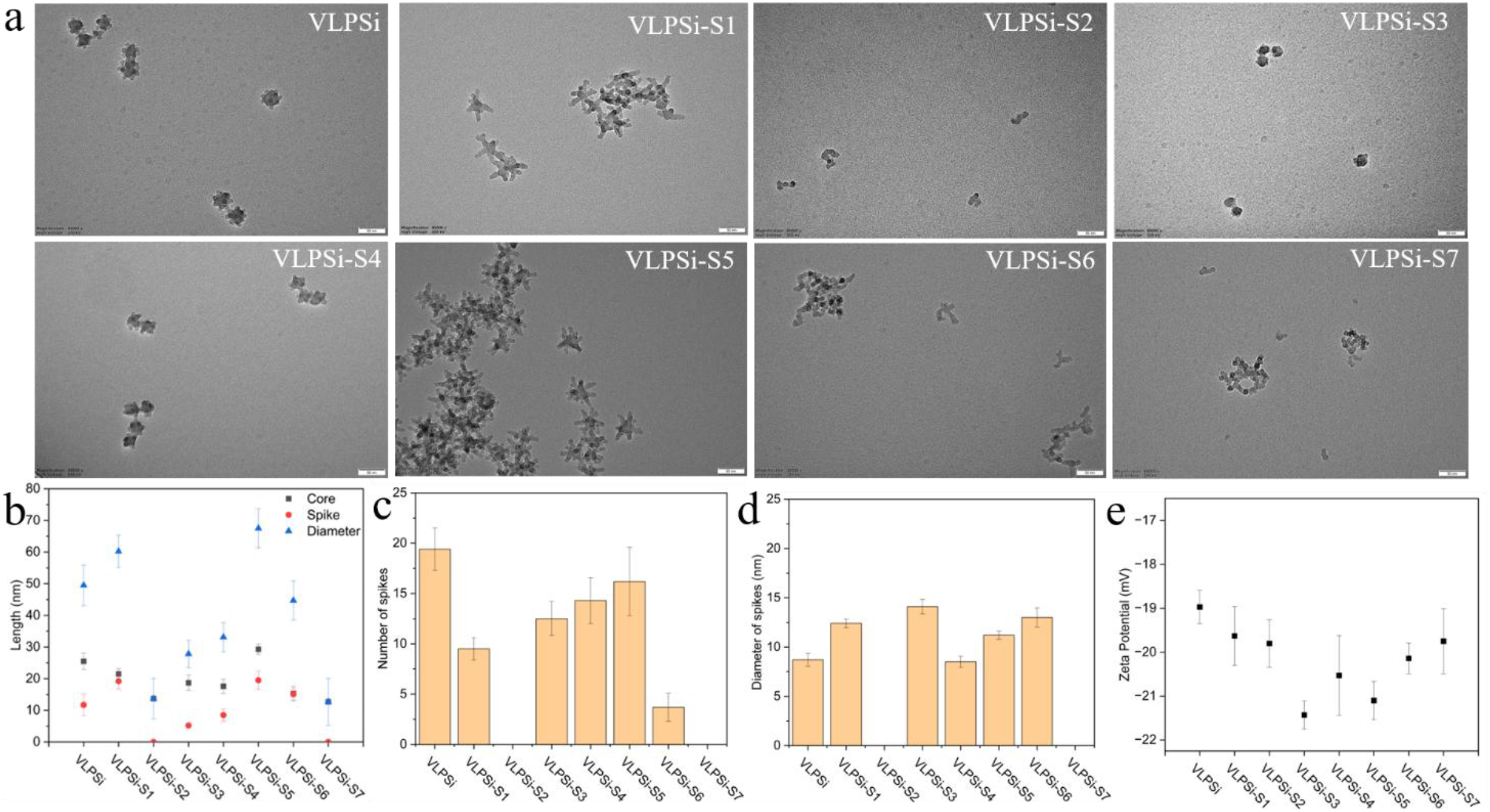
(a) TEM images of different nanoparticles. The scale bar is 50 nm. (b) Statistical data on the diameter, core size, and spike length of different nanoparticles. (c) Statistics on the number of spikes in different nanoparticles. (d) Statistics on the diameter of spikes in different nanoparticles. (e) Zeta potential of different nanoparticles.

#### 3.1.1 BTES affects the size of the pores and core of nanoparticles, ultimately influencing the morphology and structure of nanoparticles

We first tried adding 0.5 mL of BTES to the original reaction system and kept other reaction conditions unchanged to obtain VLPSi-S1. Compared with VLPSi, the diameter of the nanoparticles increased from 49.5 ± 6.4 nm to 60.2 ± 5.1 nm. This is because the spike length increased from 11.7 ± 3.4 nm to 19.2 ± 2.4 nm, despite a slight decrease in core size. In addition, the number of spikes also decreased from 19.4 ± 2.1 to 9.5 ± 1.1. Due to the conformation of the bulk organic spacer region and its associated steric effects, the hydrolysis and condensation rates of BTES are much lower than those of TEOS^23^. Therefore, the (CH_2_)_3_) -S-S-S-S-(CH_2_)_3_)-bonds in BTES would penetrate into the hydrophobic domains of the surfactant CTAB to enlarge the micelle and thus an increase in the pore size. The spikes grow from the pores of the nanoparticle, so the diameter of the spikes will also increase accordingly. Furthermore, the increased pore size indirectly leads to a slight decrease in the core diameter, directly resulting in a reduction in the number of spikes. This allows more unreacted silicon sources to grow on the spikes, thereby increasing their length. Subsequently, we increased the amount of BTES to further explore its impact. When the BTES content increased to 1 mL (VLPSi-S6), the diameter of the core decreased from 21.5 ± 1.7 nm to 15.3 ± 2.3 nm. Furthermore, a significant reduction in the number of nanoparticle spikes can be clearly observed. Interestingly, the length of the nanoparticle spikes also decreased from 22.5 ± 2.4 nm to 15.1 ± 1.9 nm, resulting in a significant reduction in the nanoparticle diameter. This was because the excessive addition of BTES led to an increase in the pore size, while the core diameter was still very small, insufficient to support the formation of longer spikes. This part of the results also verifies our above inference. When the BTES content was further increased to 2 mL (VLPSi-S7), we found that it was no longer possible to form a virus-like structure. Instead, many incomplete small nanoparticles were formed. This further illustrates that the excessive addition of BTES led to an increase in pore size and a decrease in core diameter, which was therefore insufficient to support the formation of complete virus-like nanoparticles.

#### 3.1.2 NaOH regulates the hydrolysis and condensation rates of TEOS and BTES, affecting the rate of BTES incorporation into CTAB, and ultimately influencing the morphology and structure of nanoparticles

Subsequently, the effect of different amounts of 0.1 M NaOH on nanoparticles was investigated. When the NaOH content decreased from 0.4 mL to 0.2 mL (VLPSi-S2), small nanorods were produced instead of forming virus-like nanoparticles. When the NaOH content increased from 0.4 mL to 0.8 mL (VLPSi-S3), the core size of the nanoparticles decreased from 21.5 ± 1.7 nm to 18.7 ± 2.4 nm. The length of the spikes also decreased from 22.5 ± 2.4 nm to 5.2 ± 0.8 nm, resulting in a significant reduction in the overall diameter. Furthermore, number of spikes also decreased significantly. Previous studies have shown that NaOH mainly acts as a catalyst in similar reactions, accelerating the hydrolysis and condensation of TEOS and BTES. In particular, it accelerates the incorporation speed of BTES into CTAB micelles, thereby expanding the pore size of CTAB micelles and thus the nanoparticles^24^. Therefore, the slow hydrolysis and condensation rates of TEOS and BTES are too low when the NaOH content is low, resulting in insufficient formation of virus-like structures within the same experimental time. Conversely, when the NaOH content is too high, the large catalytic activity significantly increases the rate of BTES incorporation into CTAB micelles, thereby increasing the diameter of spikes and leading to a significant decrease in the number of spikes per particle. In addition, the large amount of catalyst leads to rapid hydrolysis and condensation in the reaction, and insufficient silicon source content further reduces the spike length of the nanoparticles, which is only 5.2 ± 0.8 nm. This further demonstrated that excessively rapid hydrolysis and condensation rates of TEOS and BTES are not conducive to spike growth.

#### 3.1.3 The stirring speed affects the diffusion rate of TEOS/BTES from the oil phase to the aqueous phase, ultimately influencing the size of the nanoparticles and the number of spikes

Then, we investigated the effect of different stirring rates on the nanoparticle structure. When the stirring rate decreases from 300 rpm to 200 rpm (VLPSi-S5), the core size of the nanoparticles increased from 21.5 ± 1.7 nm to 29.3 ± 1.6 nm, while the length of the spikes did not change significantly. Therefore, the diameter of the nanoparticles also increased accordingly. When the stirring rate decreased from 200 rpm to 100 rpm (VLPSi-S4), the core size of the nanoparticles decreased from 29.3 ± 1.6 nm to 17.6 ± 2.2 nm, and the length of the spikes decreased from 19.5 ± 2.9 nm to 8.5 ± 1.9 nm. As a result, the diameter of the nanoparticles also decreased from 67.5 ± 6.2 nm to 33.1 ± 4.6nm. Furthermore, we observed a change in the number of nanoparticle spikes as the stirring rate changed. When the stirring rate decreased from 300 rpm to 200 rpm, the number of spikes increased significantly. When the stirring rate decreased from 200 rpm to 100 rpm, the number of spikes did not change significantly. Theoretically, stirring speed affects the diffusion rate and hydrolysis rate of TEOS and BTES from oil droplets to the aqueous phase^25^. Therefore, at a lower stirring speed (100 rpm), the diffusion speed of TEOS and BTES from cyclohexane to the aqueous phase is lower, resulting in a lower concentration of silica monomers at the oil-water interface, thus forming smaller nanoparticles. As the stirring rate increases (200 rpm), the diffusion speed of silica monomers also increases. The monomer concentration at the oil-water interface increases, and therefore the size of the nanoparticles increases accordingly. As the stirring rate increases further (300 rpm), the diffusion speed of the monomer also increases further, which in turn reduces the existence time of the monomer at the oil-water interface, resulting in slower growth of nanoparticles and a decrease in the diameter of the nanoparticles. Specifically, this is reflected in a decrease in the size of the core and a decrease in the number of spikes. Therefore, in this system, a stirring rate of 200 rpm can yield nanoparticles with the largest size and the highest number of spikes. Previous studies have also yielded conclusions consistent with our findings^26^.

#### 3.1.4 Physicochemical characterization of VLPSi and VLPSi-S

Finally, the chemical composition of VLPSi-S1 was investigated. VLPSi-S1 exhibits weak characteristic S-S absorption peaks in the 520-720 nm range compared to VLPSi^24^ (Figure S1a). VLPSi-S1 has weak C and S absorption peaks around 285 eV and 164 eV but VLPSi doesn’t have, further confirming the successful incorporation of tetrasulfide bonds into the nanoparticle (Figures S1b). Besides, the zeta potential of VLPSi was –18.97 ± 0.38 mV, and the VLPSi-S1 potential was – 19.63 ± 0.67 mV (Figure 1e). The result showed the zeta potential did not change significantly after the addition of BTES, which also indicates that the incorporation of tetrasulfide bonds does not affect the surface charge of the nanoparticles.

### 3.2. The structure of nanoparticles affects cell uptake efficiency

The physicochemical properties of nanoparticles, such as size, surface charge, morphology, and surface modification, significantly affect their interaction with the cell membrane, thereby regulating the efficiency and mechanism of nanoparticles and cargo entry into cells^27,28^. Five nanoparticles with same BTES content (0.5 mL) but different morphologies were selected for comparison with VLPSi. The nanoparticles were modified with FITC after they were aminated for confocal microscopy observation, and then their zeta potentials were measured. The zeta potential of all nanoparticles changed from negative to positive and was around +17 mV to +20 mV (Figure S2a). In the confocal experiment, VLPSi-S1 and VLPSi-S5 exhibited the highest FITC fluorescence areas, followed by VLPSi, VLPSi-S4, and VLPSi-S2, which showed no significant differences from one another (Figure 2). VLPSi-S3 has the smallest green fluorescence area (Figure S2b). Consistent with previous studies^11,29^, the longer spikes tend to result in higher cellular uptake efficiency. Interestingly, although VLPSi-S1 and VLPSi-S5 have similar spike lengths, their fluorescence results differ, indicating that the number of spikes also affects cellular uptake efficiency. Furthermore, VLPSi-S2, a special small nanoparticle without spike structure, also exhibits significant fluorescence, suggesting that smaller nanoparticles are more easily enter into cells. Other studies have also reached conclusions consistent with our findings^30^. In summary, the length and number of spikes significantly affect cellular uptake when nanoparticles have similar particle size. Longer and more numerous spikes generally make it easier for the nanoparticles to interact with the cell membrane, resulting in higher cellular uptake.

**Figure 2.**
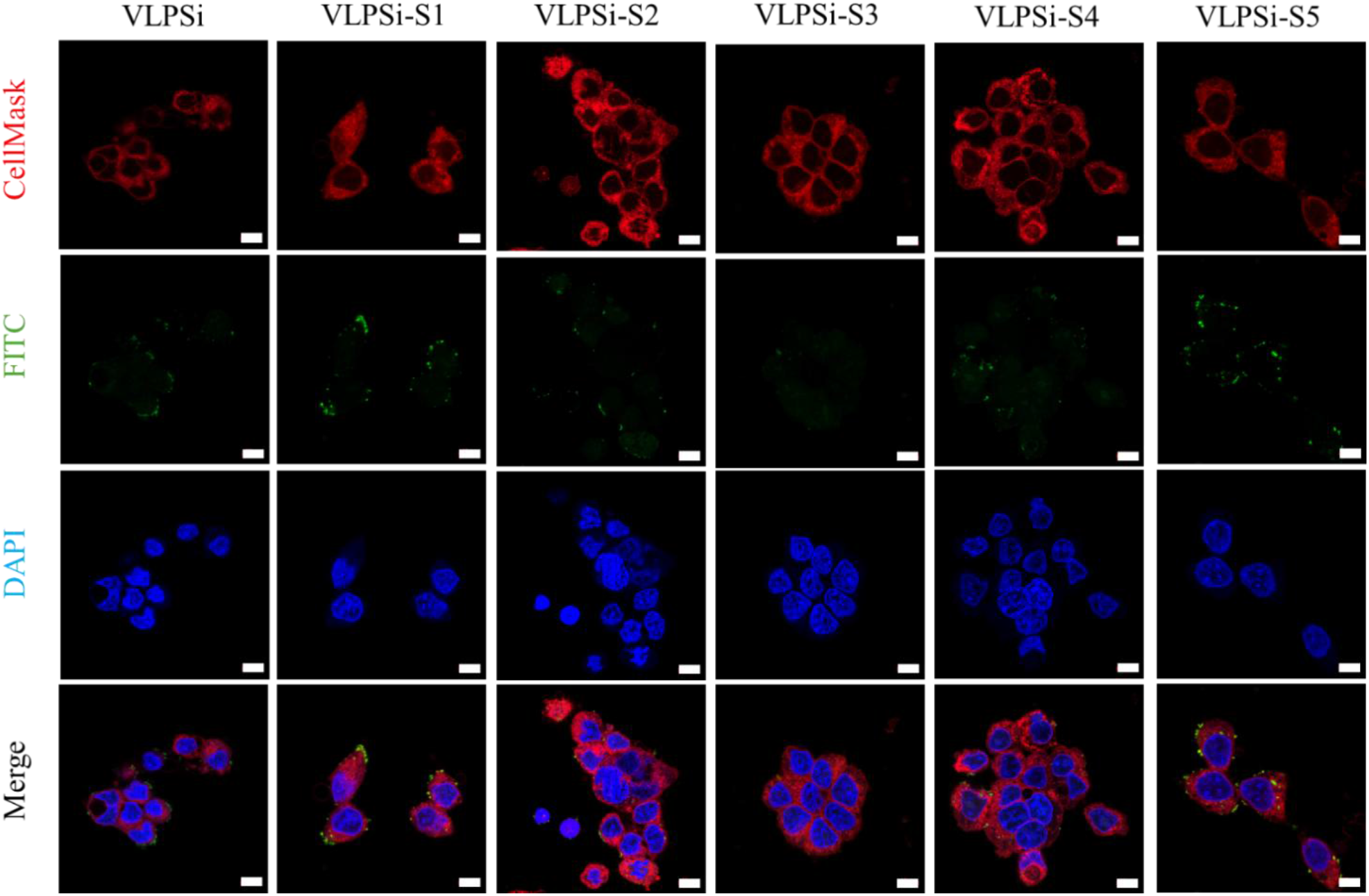
Fluorescence images of HEK-293T cells are treated with different FITC-labeled nanoparticles. Nanoparticles are labeled FITC (green), cell nucleuses are labeled DAPI (blue), and cell membranes are labeled Cell Mask (red). White scale bar: 10 μm.

### 3.3. Nanoparticles with tetrasulfide bonds have better biodegradability

To achieve effective mRNA delivery, nanocarriers should remain stable extracellularly while undergoing rapid degradation within the intracellular environment to facilitate mRNA release. Tetrasulfide bonds tend to break when exposed to intracellular high GSH. Therefore, we investigated the biodegradability of nanoparticles containing tetrasulfide bonds (VLPSi-S1) at different GSH concentrations. As shown in Figure 3, when VLPSi-S1 was in simulated environment with 5 µM GSH, it was very stable at all time points and no degradation occurred, indicating that the low GSH environment such as extracellular/plasma was insufficient to cause VLPSi-S1 degradation. In contrast, progressive degradation was observed at 5 mM GSH. The particle edges became blurred after 8 h, followed by substantial dissociation of the surface structures at 16 h and the appearance of fragmented structures at 24 h. Degradation was further accelerated at 10 mM GSH, with fragmented structures already apparent at 8 h. The virus-like morphology was largely lost by 16 h, and the particles were extensively degraded by 24 h. In contrast, VLPSi maintained its morphology and particle size under all tested GSH concentrations (Figure S3). In summary, VLPSi-S1 possess good GSH-responsive intracellular biodegradability while remaining generally stable in extracellular/plasma environments.

**Figure 3.**
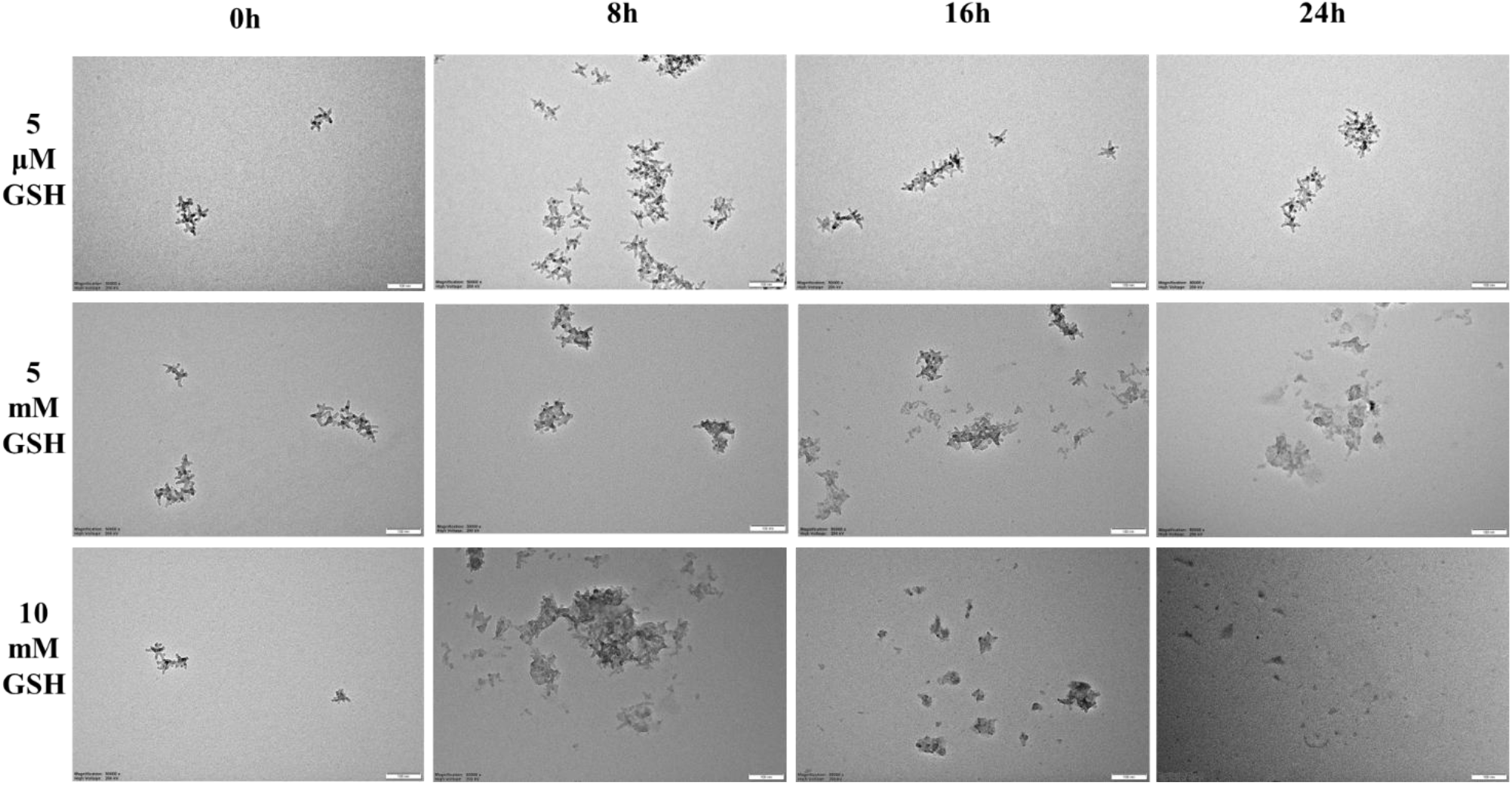
TEM images of VLPSi-S1 at different time points under simulated environments with different GSH concentrations. 5 μM, 5 mM, 10 mM GSH correspond to the simulated concentrations of GSH in extracellular/plasma environment, intracellular environment of healthy cells, and the intracellular environment of tumor cells, respectively. The scale bar in the figure is 100 nm.

### 3.4. Different nanoparticles all have good biocompatibility

The safety of nanoparticles is also a crucial prerequisite for biological research^31^. Therefore, we investigated the cytotoxicity of various nanoparticles in HEK-293T cells. For our experimental purposes, nanoparticles were used to load mRNA for transfection experiments, and the required nanoparticle concentration was typically about 10 μg/mL. When the concentration of nanoparticles was 10 μg/mL, the cell viability of all nanoparticles was greater than 95% (Figure S4). In summary, these six nanoparticles are highly safe for cells at a concentration of 10 μg/mL and can be used for further research on subsequent mRNA transfection.

### 3.5. Different nanoparticles get different mRNA transfection effects

To evaluate the mRNA delivery, mCherry mRNA was initially complexed with ionizable lipid ALC-0315 and the helper lipid DOPC (ALC-0315/DOPC) and subsequently incorporated with the nanoparticles. Transfection efficiency was quantified by flow cytometry using the gating strategy (Figure S5a). The mCherry mRNA expressions of VLPSi-S1, VLPSi-S2, VLPSi-S4, and VLPSi-S5 were significantly higher compared to VLPSi (Figure 4a and 4b), whereas no significant difference was observed between VLPSi-S3 and VLPSi. Overall, the transfection trend was consistent with the cellular uptake results, supporting a contribution of nanoparticle internalization to mRNA delivery. Interestingly, although VLPSi exhibited a cellular uptake efficiency comparable to that of VLPSi-S4, its mCherry mRNA % was significantly lower than that of VLPSi-S4. In summary, most nanoparticles incorporating tetrasulfide bonds exhibited higher transfection efficiency than VLPSi. Among them, only VLPSi-S3 showed a comparable effect to VLPSi, which can be attributed to its relatively lower cellular uptake efficiency. The median fluorescence intensity (mCherry mRNA MFI) further revealed differences in mRNA expression among the groups (Figure 4c). All tetrasulfide bond-containing nanoparticles exhibited significantly higher than VLPSi. Among them, VLPSi-S5 exhibited markedly higher MFI than all other groups, followed by VLPSi-S1, VLPSi-S4, VLPSi-S2, and VLPSi-S3, while VLPSi showed the lowest fluorescence intensity. These results were consistent with the mCherry mRNA % but provided greater separation among the groups. To further validate these findings, firefly luciferase (Luc)-encoding mRNA was subsequently employed. The fluorescence intensity and statistical differences among the experimental groups were fully consistent with those observed in Figure 4c, further confirming the reliability of our findings (Figure S5b). In summary, the incorporation of tetrasulfide bonds effectively enhances transfection efficiency. Specifically, these bonds likely facilitate the accelerated degradation of nanocarriers and the subsequent intracellular release of mRNA, thereby promoting transfection and supporting our initial hypothesis. In addition, previous studies have reported that the high intracellular GSH environment can hinder transfection28. Tetrasulfide bonds can undergo redox reactions with intracellular GSH, which may reduce GSH levels and thereby alleviate its inhibitory effect on mRNA translation. This process can further enhance transfection efficiency. Consequently, VLPSi-S5 demonstrates the highest transfection performance, which can be attributed to the incorporation of tetrasulfide bonds and its more and longer spiked structure.

**Figure 4.**
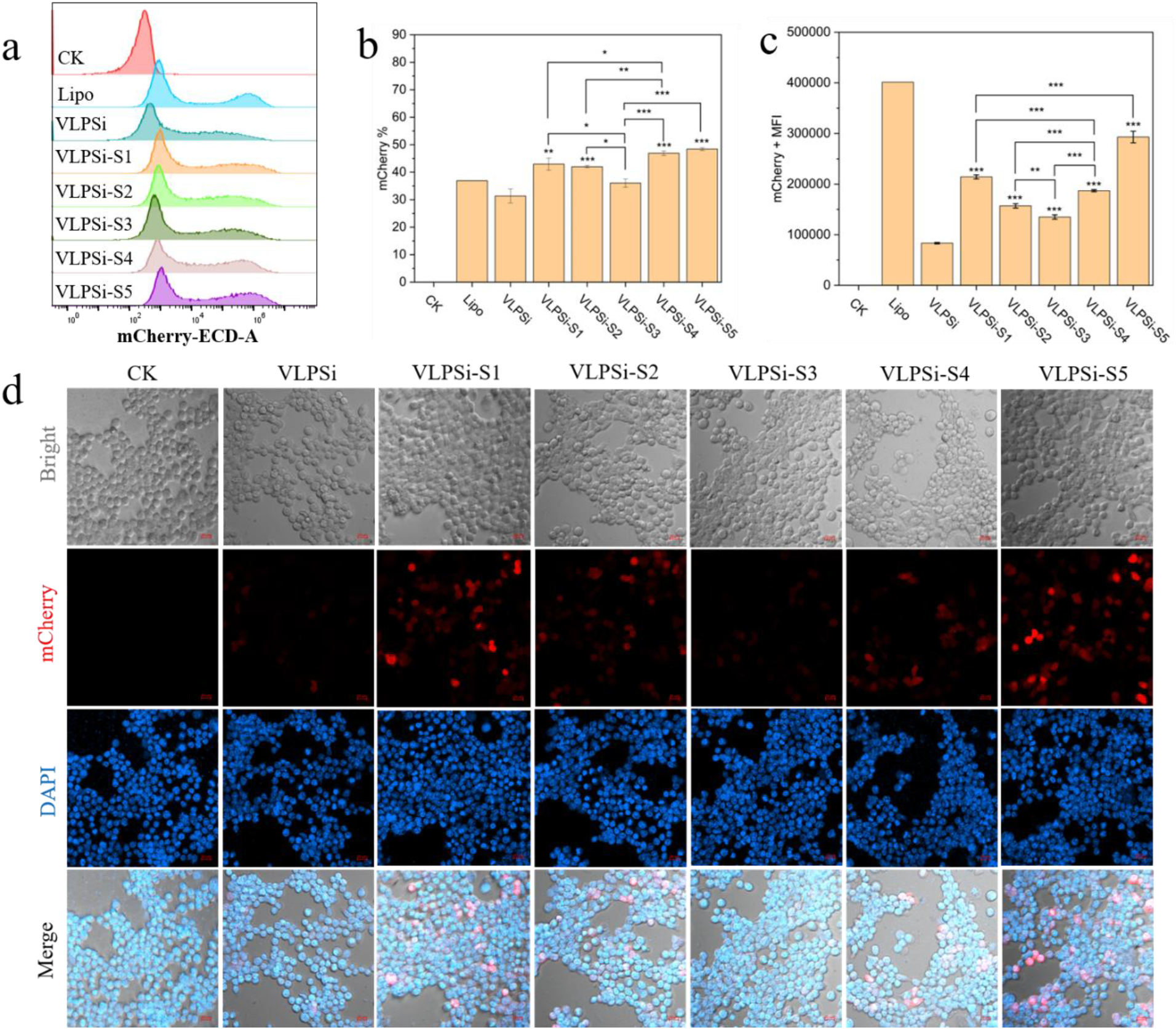
(a) Flow cytometry histogram of mCherry mRNA expression on HEK-293T cells after transfection with control check (CK), Lipo, and different nanoparticles groups. (b) mCherry mRNA % result and (c) Mean MFI value of cells transfected with different nanoparticles loaded with lipids and mCherry mRNA. (d) Transfection fluorescence images of cells transfected by different nanoparticles loaded with mCherry mRNA. Cells, nucleus, mCherry mRNA expression are shown in bright field, blue (DAPI), and red channels, respectively. Scale bar is 20 micrometers.

To further validate the flow cytometry results, confocal microscopy was employed to visualize mCherry mRNA expression and assess fluorescence intensity (Figure 4d). Identical regions of interest with equal areas were selected from each image to ensure consistency in the analysis of fluorescence area and intensity. The fluorescence area of VLPSi-S5 was significantly greater than that of all other groups. The fluorescence areas of VLPSi-S1, VLPSi-S2, and VLPSi-S4 were comparable and markedly higher than VLPSi and VLPSi-S3 (Figure S6a). Similarly, the mCherry mRNA fluorescence intensity of VLPSi-S5 was the highest among all experimental groups, followed by VLPSi-S1, VLPSi-S4, VLPSi-S2, VLPSi-S3, and VLPSi (Figure S6b). These trends were consistent with the mCherry mRNA % and MFI measured by flow cytometry, further validating our conclusion above.

### 3.6. Nanoparticles loaded with different lipids enhanced the mRNA delivery efficiency

To assess the versatility of the nanoparticle platform, we further evaluated different lipid formulations using the cationic lipids DOTAP and DOTMA to replace ALC-0315, and VLPSi-S5 was chosen as carrier. Their transfection performance was compared with that of lipid – mRNA complexes without nanoparticles. The gating strategy is presented in Figure S7a. Regardless of the type of lipids, the addition of nanoparticles can greatly improve the transfection effect of mCherry mRNA, and there is a significant difference compared with the transfection effect of lipid – mRNA complex only (Figure 5a and Figure 5b). Subsequently, the overall trend of mCherry mRNA MFI is the same as that of mCherry mRNA % (Figure 5c). These findings were further validated using Luc mRNA, which produced consistent fluorescence and statistical trends (Figure S7b). In summary, the nanoparticles we synthesized can effectively load different lipids and mRNA for transfection, greatly improving the mRNA transfection efficiency. This indicates that the nanoparticles can serve as a universal carrier to assist different lipids and mRNAs in transfection.

**Figure 5.**
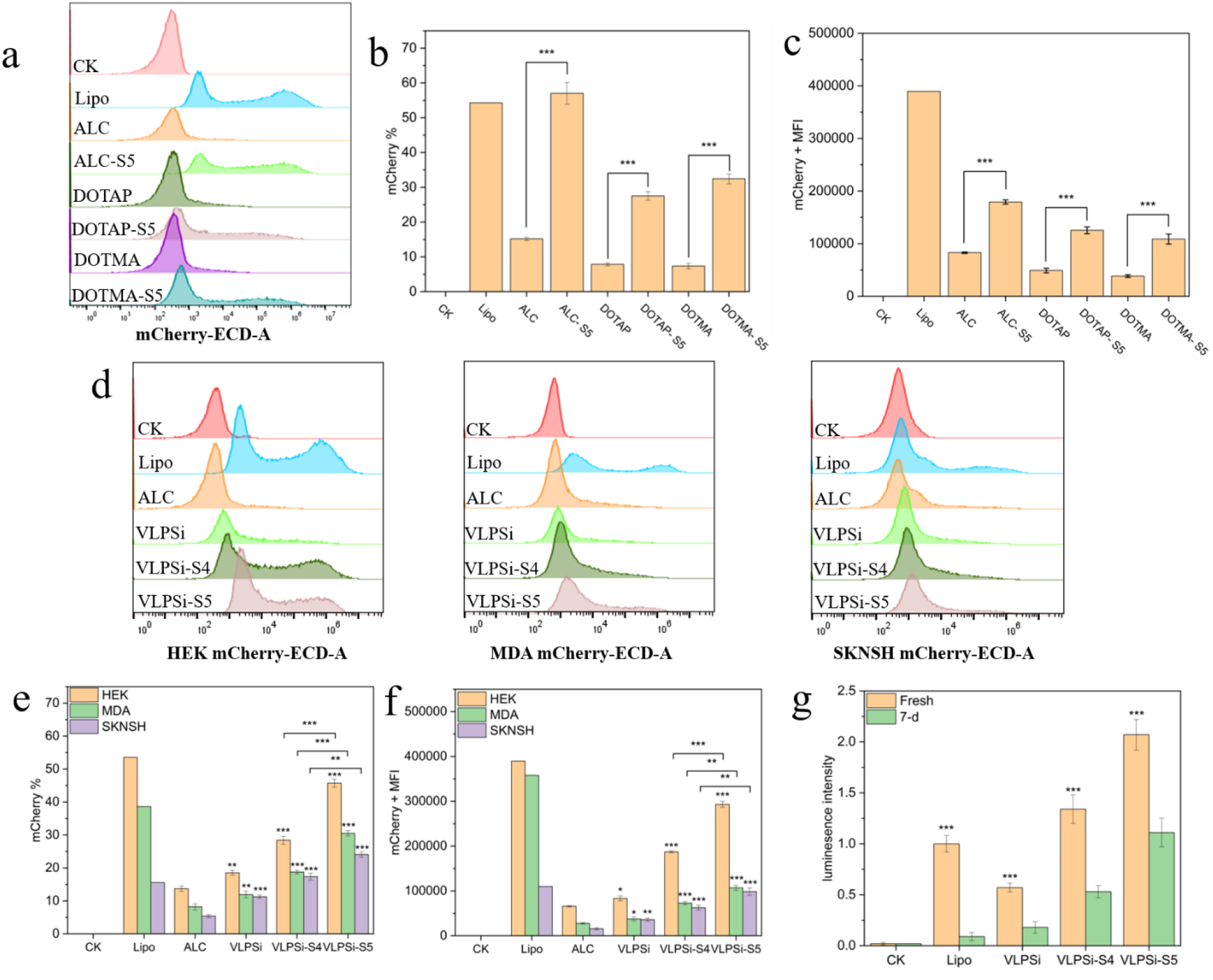
(a) Flow cytometry histogram of mCherry mRNA expression on HEK-293T cells after transfection with control check (CK), Lipo, and different lipids groups. (b) mCherry mRNA % result and (c) Mean MFI value of cells transfected with nanoparticles loaded with different lipids and mCherry mRNA. (d) Flow cytometry histogram of mCherry mRNA expression on different cells after transfection with control check (CK), Lipo, and different lipids groups. (e) mCherry mRNA % result and (f) Mean MFI value of different cells transfected with nanoparticles loaded with lipids and mCherry mRNA. (g) Analysis of the transfection effect of different nanoparticle loaded lipids and Luc mRNA with fresh preparation and storage for 7 days.

### 3.7. Efficient mRNA delivery across different cell lines

To further evaluate the transfection performance of nanoparticles, we determine transfection efficacy on MDA and SKNSH cell lines. Typically, SKNSH cells are recognized as difficult to transfect^32,33^. MDA cells have higher GSH concentration in its intracellular environment than normal cells, thus allowing for further comparative studies^34^. VLPSi-S4 exhibits similar spike length and cellular uptake efficiency to VLPSi. The two differ only in their internal chemical structures, making them suitable candidates for comparison. Therefore, we selected VLPSi, VLPSi-S4, and VLPSi-S5 as nanocarriers for this part of research. Different nanoparticles were used to load ALC-0315/DOPC and mCherry mRNA then transfected on different cell lines. Lipo and ALC-0315/DOPC-mCherry mRNA were used as positive controls. The gating settings are shown in Figure S7. Regardless of the cell line, the transfection effect of VLPSi-S5 was significantly higher than that of VLPSi-S4, and the transfection effect of VLPSi-S4 was significantly higher than that of VLPSi (Figure 5d and Figure 5e). MFI showed the same overall trend (Figure 5f). Notably, VLPSi-S5 achieved mCherry mRNA % of 24.08 ± 0.84% in SKNSH and 30.5 ± 0.82% in MDA cells, indicating significant transfection in these relatively difficult-to-transfect cell models.^35,36^. The results were further validated using Luc mRNA, which showed consistent fluorescence and statistical trends (Figure S8b). In summary, the nanoparticles we synthesized, especially VLPSi-S5, can achieve highly efficient transfection in different cell lines after loading lipids and mRNA.

### 3.8. VLPSi-S5 significantly enhanced the stability of mRNA

Because mRNA is easily degraded, storage stability is an important consideration for its wider application.^37,38^. We selected the three nanoparticles used above to further study their stability after loading lipids and mRNA. The three nanoparticles were formulated with ALC-0315/DOPC and Luc mRNA and stored at the 4°C for 7 days. Lipo-Luc mRNA complex was subjected in the same condition as a control. Then the fresh samples were prepared and transfected into cells together with the old samples (Figure 5g). Lipo exhibited high transfection efficiency when freshly prepared but showed a marked decrease after 7 days at 4 °C. In contrast, our nanoparticle loaded lipids and mRNA maintained some effectiveness after 7 days, corresponding to approximately 31%, 40%, and 54% of the freshly prepared transfection efficiency for VLPSi, VLPSi-S4, and VLPSi-S5, respectively. In summary, our synthesized nanoparticles significantly improve the storage stability of mRNA, especially VLPSi-S5. We believe this is due to its unique spiked structure, which protects the loaded mRNA and lipids from external contact, thus stabilizing the mRNA structure.

## 4. Conclusions

We developed tetrasulfide-incorporated mesoporous silica nanoparticles and explored their potential as a carrier for mRNA delivery. First, tetrasulfide bonds were incorporated into previously synthesized VLPSi. By adjusting different experimental parameters, nanoparticles with different morphologies were synthesized. Then, the effects of different structures and chemical compositions on mRNA transfection efficiency were compared. The results showed that tetrasulfide incorporation significantly influenced the formation of mesoporous silica nanoparticles. By optimizing reaction conditions, virus-like mesoporous silica nanoparticles with uniform morphology and intact structure could also be obtained, clarifying the regulatory role of key factors in the virus-like structure.

Furthermore, among mesoporous silica nanoparticles with different structures and chemical compositions, virus-shaped nanoparticles exhibited the highest cellular uptake efficiency and low cytotoxicity. The length and number of spike structures also significantly affected cellular uptake efficiency and transfection efficacy. Compared to nanoparticles without tetrasulfide bonds, nanoparticles containing tetrasulfide bonds exhibit rapid and responsive degradation within cells, promoting mRNA release from the carrier while simultaneously reducing intracellular GSH concentration. This ultimately promotes mRNA translation within cells and further enhances transfection. In summary, VLPSi-S6 achieves the best transfection efficiency due to its longest and most numerous spikes, as well as the presence of tetrasulfide bonds. Therefore, this study provides a novel strategy for constructing mesoporous silica-based nucleic acid delivery systems with controllable degradation characteristics, laying the foundation for their application in gene therapy and mRNA delivery.

## CRediT authorship contribution statement

**Zhe Cao:** Writing – original draft, Validation, Formal analysis, Data curation, Investigation, Methodology, Writing – review & editing. **Silja Saarela:** Writing – review & editing, Visualization, Methodology, Formal analysis, Data curation, Conceptualization. **Jiajia Wang:** Writing – review & editing, Visualization, Methodology, Formal analysis, Data curation, Conceptualization. **Minna Sivonen:** Writing – review & editing, Visualization, Methodology, Formal analysis, Data curation, Conceptualization. **Emilia Happonen:** Methodology, Formal analysis, Data curation. **Jussi Hepojoki:** Writing – review & editing, Methodology, Supervision. **Hiromi Muramatsu:** Methodology, Formal analysis. **Pardi Norbert:** Writing – review & editing, Methodology, Supervision. **Vesa-Pekka Lehto:** Writing – review & editing, Supervision, Resources, Project administration, Conceptualization. **Wujun Xu:** Writing – review & editing, Supervision, Project administration, Funding acquisition.

## Acknowledgments

This work was supported by Research Council of Finland (Grant no. 356056) and China Scholarship Council. SIB Labs and Cell & the Tissue Imaging Unit at the University of Eastern Finland were acknowledged for technical support.

## Data Availability

Data will be made available on request.

