## Supplementary Materials for "Engineering morphology of tetrasulfide-incorporated mesoporous silica nanoparticles for enhanced mRNA delivery"

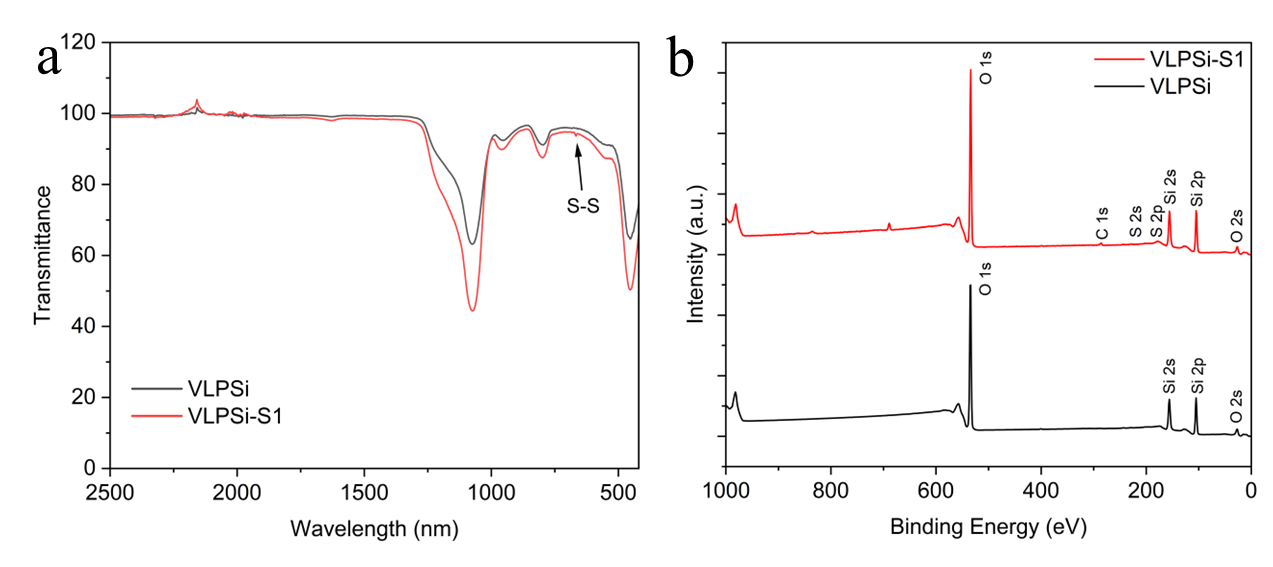

Figure S1. (a) FTIR images of VLPSi and VLPSi-S1. (b) XPS image of VLPSi and VLPSi-S1.

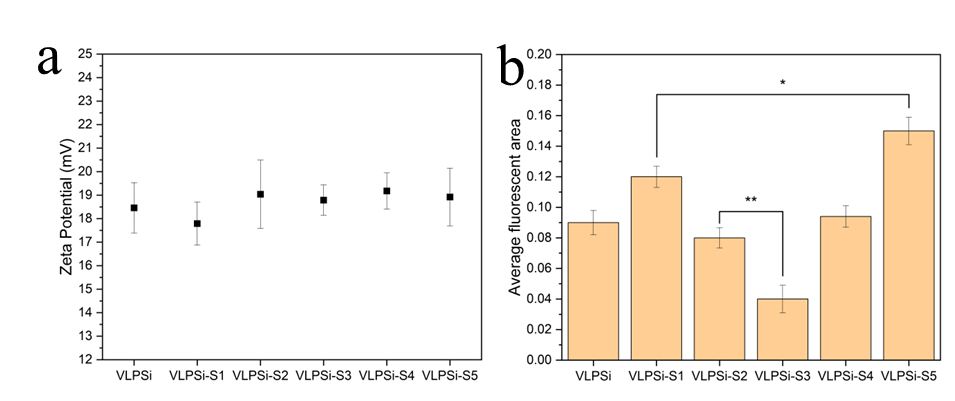

Figure S2. (a) Zeta potential of amino groups modified with different nanoparticles. (b) Average fluorescence intensity of FITC in each experimental group.

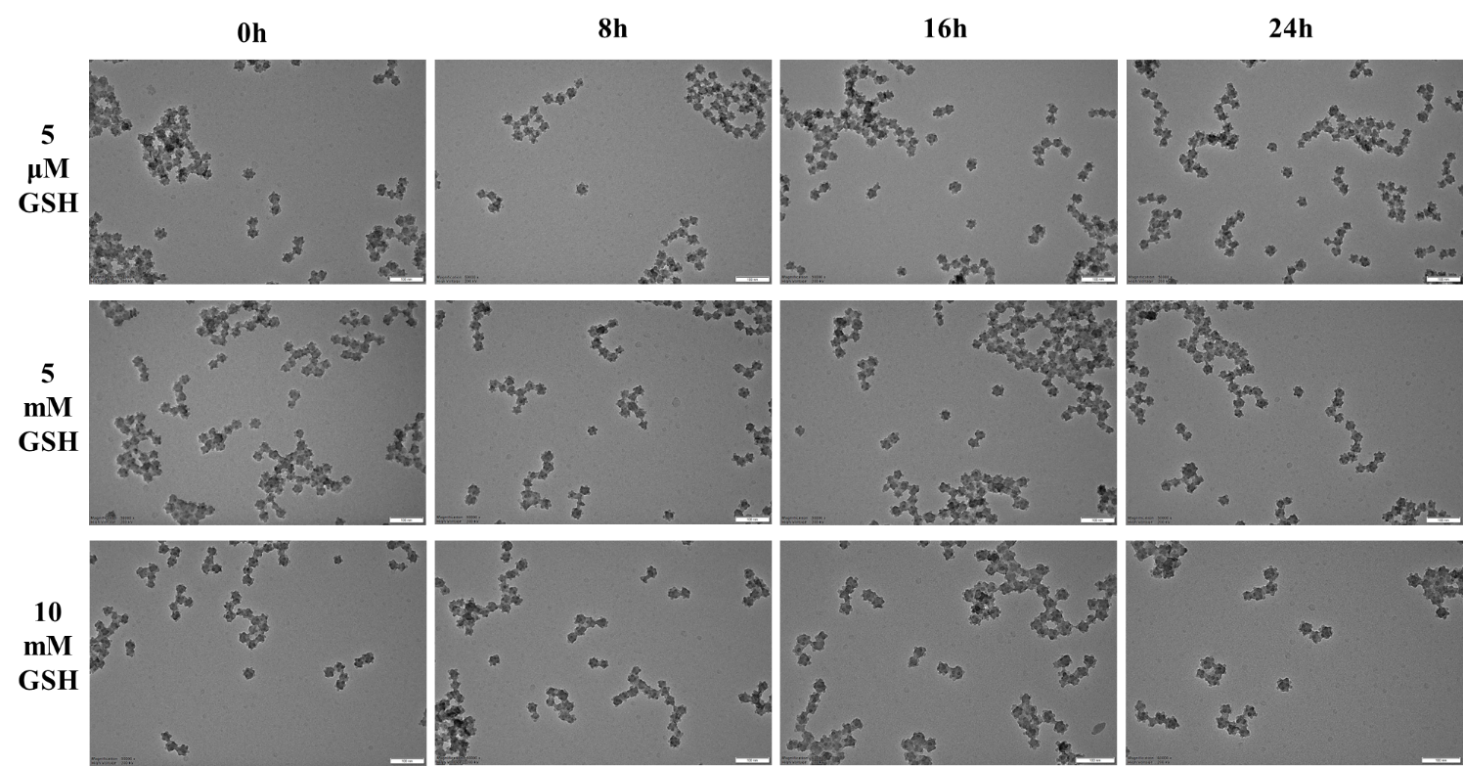

Figure S3. TEM images of VLPSi-S1 at different time points under simulated environments with different GSH concentrations. The scale bar is 100 nm.

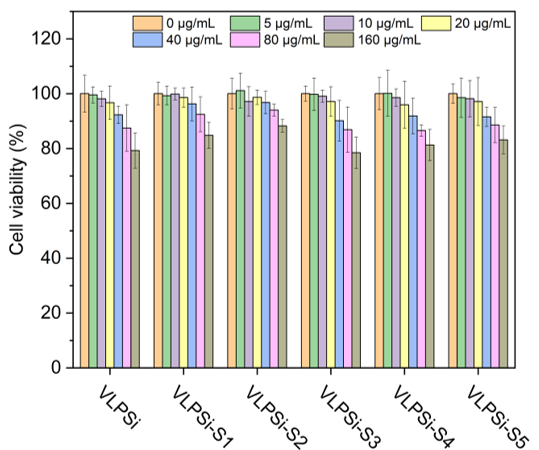

Figure S4. HEK-293T cell viability results after treatment with different nanoparticles.

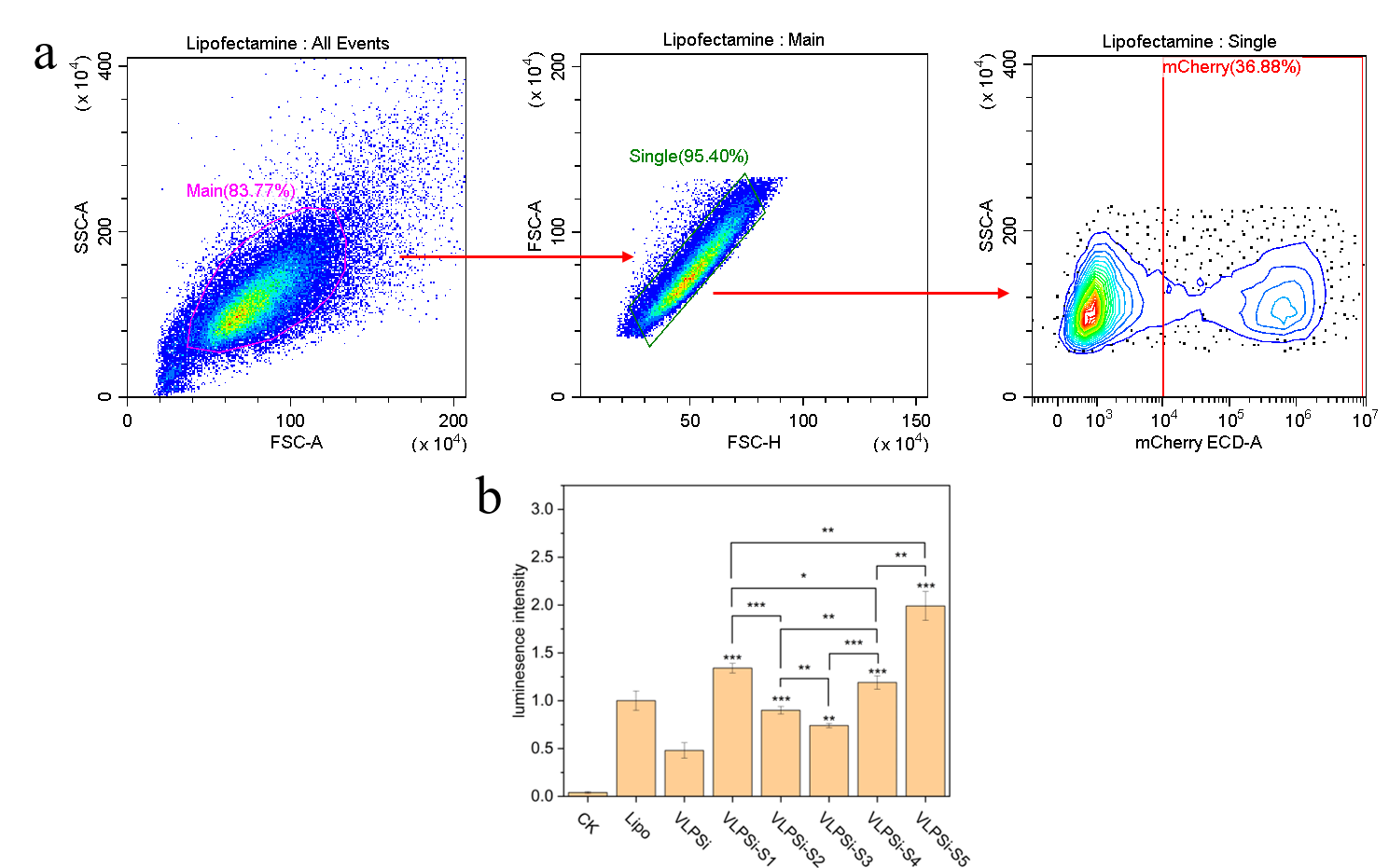

Figure S5. (a) Flow cytometry gating strategies. The data was processed using FlowJo. Taking Lipo as an example, all events were first preliminarily screened based on FSC-A and SSC-A to delineate the main cell population (83.77%). Subsequently, cell aggregates were removed from the main cell population using FSC-H and FSC-A to perform single-cell population screening (95.40%). Finally, transfection efficiency was assessed in the single-cell population based on mCherry mRNA fluorescence intensity (mCherry ECD-A) (positive cell rate was 36.88%). (b) Mean luminescence intensity results of cells transfected with nanoparticles loaded with Luc mRNA.

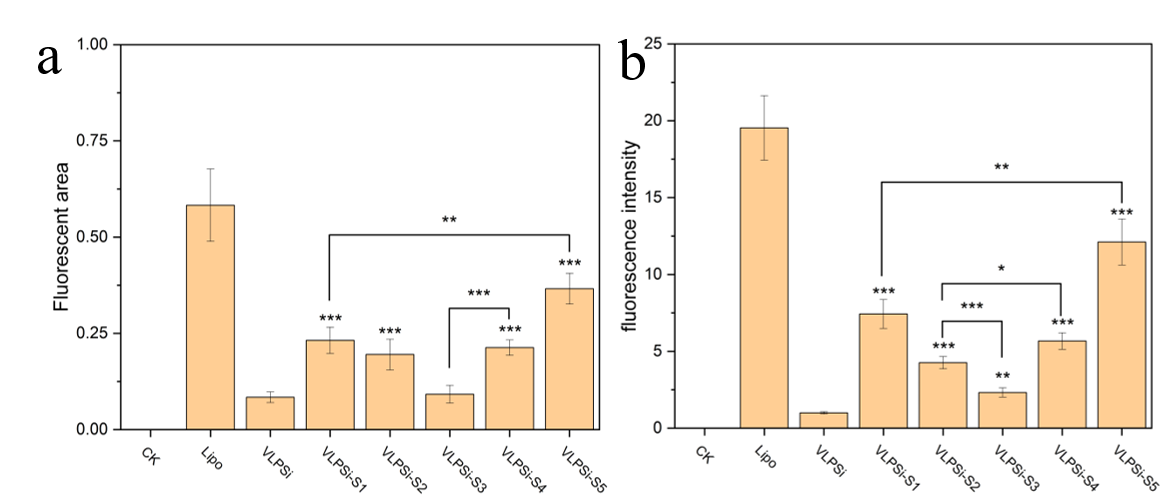

Figure S6. (a) statistics of mCherry mRNA red fluorescent area in each experimental group. (b) statistics of mCherry mRNA red fluorescence intensity in each experimental group.

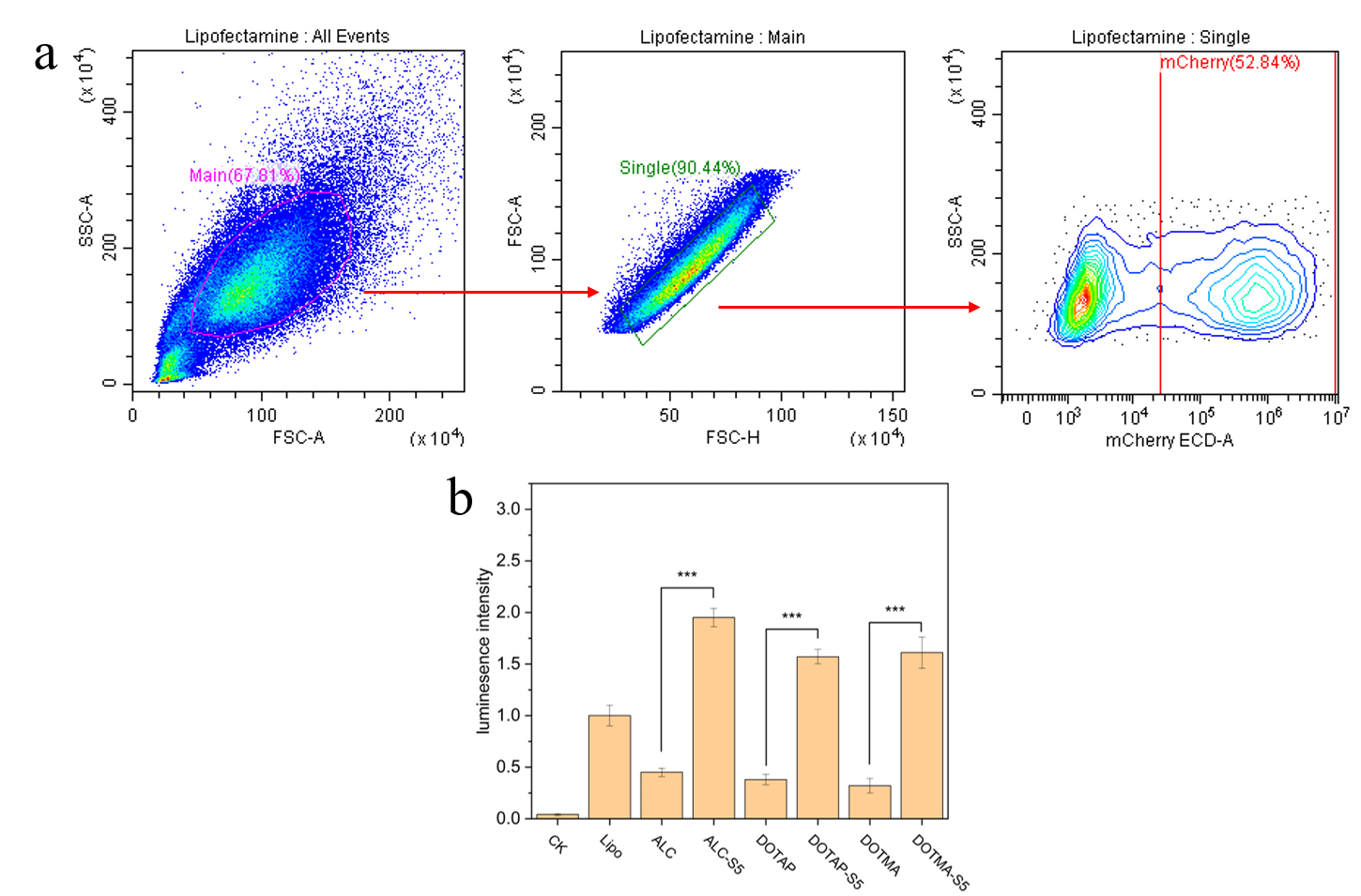

Figure S7. (a) Flow cytometry gating strategies. The data was processed using FlowJo. Taking Lipo as an example, analytical methods are same as Figure S5. (b) Mean luminescence intensity results of cells transfected with nanoparticles loaded with Luc mRNA and different lipids.

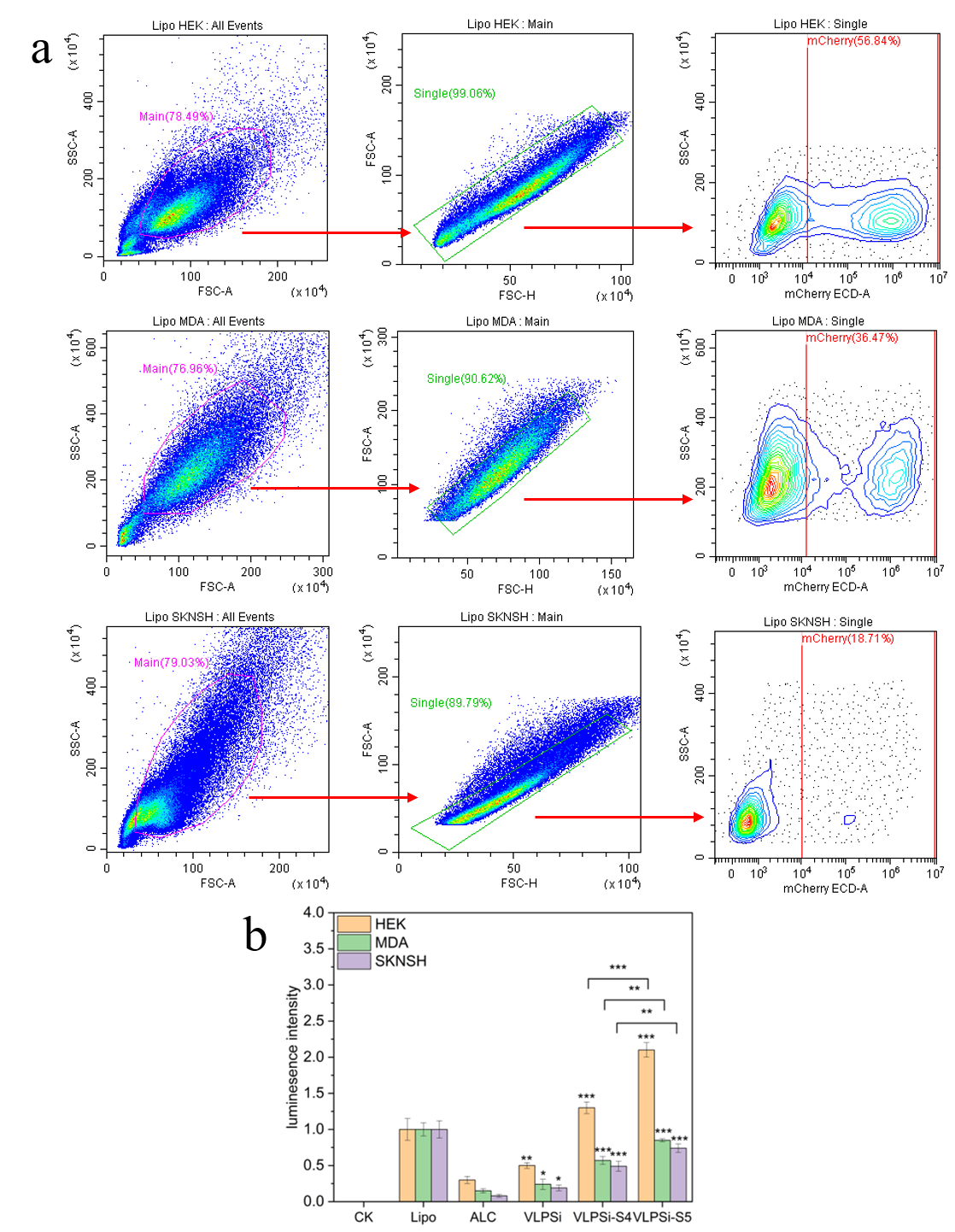

Figure S8. (a) Flow cytometry gating strategies of different cell lines. The data were processed using FlowJo. Taking Lipo as an example, the analysis methods for different cell lines are same as in Figure S4. (b) Mean luminescence intensity results of different cells transfected with nanoparticles loaded with Luc mRNA and lipids.
